# EXOSC10-mediated pre-tRNA surveillance safeguards neuron survival

**DOI:** 10.1101/2025.06.19.660564

**Authors:** Yoshinori Nishimoto, Xin Yang, Richard I. Gregory

## Abstract

tRNA quality control pathways have been identified in yeast, whereby aberrant and hypomodified mature tRNAs are targeted for 5’-3’ degradation by the rapid tRNA decay (RTD) pathway involving the Xrn1 and Rat1/Xrn2 exonucleases, whereas aberrant precursor tRNAs (pre-tRNAs) are targeted for 3’-5’ degradation by the nuclear surveillance pathway involving the RNA Exosome. However, the pathways controlling tRNA and pre-tRNA degradation in mammals have not yet been defined and the relevance of pre-tRNA surveillance pathways for normal cell physiology remains largely unknown. The RNA Exosome comprises a core of nine non-catalytic subunits (EXOSC1-9) to which the distinct, DIS3 and EXOSC10, 3’-5’ exonucleases associate. Here we find that EXOSC10 deficiency leads to accumulation of unspliced precursor tRNAs (pre-tRNAs) in mouse embryonic stem cells (ESCs) and is required for pre-tRNA decay in biochemical assays. Pre-tRNA overexpression causes diminished motor neuronal survival in a mouse ESC differentiation model. Our results identify a pre-tRNA decay pathway that links Exosome deficiency with neuron survival and provides insight into possible pathological mechanisms underlying human neurodevelopmental disorders caused by mutations in Exosome subunits and genes involved in tRNA biogenesis.

- Exosc10 is required for pre-tRNA surveillance in cells and degradation in biochemical assays
- Pre-tRNA expression inhibits survival of neurons
- Pre-tRNA decay pathway links exosome deficiency with neurodevelopmental disorders

## INTRODUCTION

tRNA biogenesis involves a complex series of events that begins with their transcription by RNA Polymerase III (Pol-III) and involves cleavage of the 5’ leader sequence by the RNase P complex, cleavage of the 3’ trailer sequence by RNase Z, CCA addition to 3’-end by tRNA-nucleotidyltransferase 1 (TRNT1), and for some tRNAs, removal of an intron (Phizicky and Hopper 2023, Schultz and Kothe 2024). A subset of 28 tRNA genes from the amongst the 429 high-confidence annotated set of tRNA genes in the human genome contain a short (∼20 nucleotide) intronic sequence that is removed by the tRNA-splicing complex during tRNA maturation (Chan and Lowe 2016). In humans, this list of intron-containing tRNAs includes 5 (out of 6) tRNA-Arg-TCT genes, 5 (out of 6) tRNA-Leu-CAA genes, 5 (out of 5) tRNA-Ile-TAT genes, and 13 (out of 13) tRNA-Tyr-GTA genes (Chan and Lowe 2016). Precursor tRNA (pre-tRNA) splicing involves the tRNA splicing endonuclease (TSEN) complex comprising four core subunits, TSEN2, TSEN34, TSEN15, and TSEN54 (Hayne, Butay et al. 2023, Yuan, Han et al. 2023, Zhang, Yang et al. 2023). In addition to these orchestrated processing events, mature tRNAs are subject to an extensive array of chemical modifications, with a typical tRNA having an average of ∼13 different RNA modifications from a pool of ∼50 modifications known to occur on tRNAs (Suzuki 2021). These modifications occur in the anticodon loop to influence mRNA translation fidelity and/or efficiency through codon-anticodon recognition and occur elsewhere in the tRNA to effect tRNA structure, function, and stability (Suzuki 2021, Schultz and Kothe 2024). Moreover, mature tRNA levels can be controlled post-transcriptionally in response to cellular stress, for example by widespread tRNA cleavage by Angiogenin, SLFN11-mediated specific cleavage of tRNA-Leu-TAA in response to DNA damage, or SAMD9/9L-mediated cleavage of phenylalanine tRNA in response to poxvirus infections (Yamasaki, Ivanov et al. 2009, Li, Kao et al. 2018, Zhang, Yang et al. 2023, Elder, Papadopoulos et al. 2024).

Work in yeast has revealed the rapid tRNA decay pathway (RTD), that ensures the fidelity of mature tRNAs, serving as a quality control step by degrading hypomodified tRNAs that lack one or more chemical modifications or with destabilizing mutations (Alexandrov, Chernyakov et al. 2006). RTD requires the 5′–3′ exonucleases Rat1 in the nucleus, and Xrn1 in the cytoplasm (Chernyakov, Whipple et al. 2008, Phizicky and Hopper 2023). Furthermore, a pre-tRNA nuclear surveillance pathway has also been described in yeast involving the TRAMP complex and the RNA Exosome that degrades unspliced, hypomodified, and/or misfolded pre-tRNAs (Alexandrov, Chernyakov et al. 2006). The RNA Exosome is involved in the processing or decay of a wide variety of different cellular RNAs (Mitchell, Petfalski et al. 1997). It comprises a core of nine non-catalytic subunits (EXOSC1-9) to which the distinct, DIS3 and EXOSC10, 3’-5’ exonucleases associate (Makino, Baumgartner et al. 2013, Januszyk and Lima 2014, Wasmuth, Januszyk et al. 2014). Transcriptome-wide analysis of Exosome targets in *S. cerevisiae* suggested that RRP44 (ortholog of mammalian DIS3) and Rrp6 (ortholog of mammalian EXOSC10) may target mature tRNA and/or pre-tRNA, as well as many other types of cellular RNAs (Schneider, Kudla et al. 2012). However, the relevance of these tRNA quality control pathways in mammalian cells remains largely unknown.

Pontocerebellar hypoplasia (PCH) is an autosomal recessive neurological disorder characterized by atrophy of the cerebellum and pons, and progressive microcephaly (Namavar, Barth et al. 2011). Among at least ten PCH subtypes, PCH type 1 (PCH1) is characterized cerebellar atrophy and defects in the motor neurons in the anterior horn of the spinal cord, leading to devastating motor neurodegeneration and muscle wasting lethally during infancy (OMIM #614678)(Goutieres, Aicardi et al. 1977). Genes mutated in several PCH subtypes are known, including *TSEN54* (tRNA splicing endonuclease 54), *TSEN34* (tRNA splicing endonuclease 34), *TSEN2* (tRNA splicing endonuclease 2), *RARS2* (arginyl-tRNA synthetase 2, mitochondrial)*, VRK1* (vaccinia-related kinase 1), and CLP1 (cleavage and polyadenylation factor I subunit 1)(Alexandrov, Chernyakov et al. 2006, Budde, Namavar et al. 2008, Renbaum, Kellerman et al. 2009, Agamy, Ben Zeev et al. 2010, Namavar, Barth et al. 2011, Burglen, Chantot-Bastaraud et al. 2012, Akizu, Cantagrel et al. 2013, Karaca, Weitzer et al. 2014). The *EXOSC3* (exosome component 3) gene was identified as the causative gene of the PCH1B-subtype (Wan, Yourshaw et al. 2012). Interestingly, mutations in another core exosome subunit, *EXOSC8* (exosome component 8) have been linked with the PCH1C subtype (Boczonadi, Muller et al. 2014). The RNA exosome complex participates in a multitude of cellular RNA processing and degradation events. Despite genetic evidence linking EXOSC3 and EXOSC8 with PCH1, the molecular and cellular links with disease remain unknown.

Several PCH-associated genes have important roles in regulating tRNA metabolism (Blaze and Akbarian 2022). This indicates that misregulation of tRNA maturation might have a critical impact on neuron differentiation and/or survival. Supporting this hypothesis, additional tRNA processing factors including the CLP1 (a kinase required for tRNA splicing) and Angiogenin (*ANG*, a stress-induced ribonuclease that cleaves tRNAs), were also reported to be associated with motor neuron dysfunction, for example, *CLP1* mutations have been reported to cause progressive spinal motor neuron loss in a kinase-dead mouse model and in human PCH10 patients (Agamy, Ben Zeev et al. 2010, Karaca, Weitzer et al. 2014, Schaffer, Eggens et al. 2014). The loss of function mutations in *ANG* were reported in amyotrophic lateral sclerosis - another motor neuron disease (Greenway, Andersen et al. 2006). ANG regulates the cleavage of tRNA (transfer RNA) into smaller fragments termed as tiRNA in the cytoplasm, which affects protein translation to promote cell survival (Emara, Ivanov et al. 2010). Altogether these findings raise the possibility that altered tRNA biogenesis and/or decay might be a common pathway contributing to PCH pathology, yet how the Exosome might be related to this pathway has not been addressed.

In this study, we explored the possible effects of Exosome deficiency on tRNA metabolism. We observed a specific accumulation of unspliced pre-tRNAs in EXOSC10-knockdown ESCs. Accordingly, we find in biochemical assays a requirement of the EXOSC10 for pre-tRNA decay. Finally, using an *in vitro* ESC differentiation model we demonstrate a pathological role of pre-tRNAs in inducing motor neuron death. Our study provides insight into how Exosome deficiency and pre-tRNA accumulation can contribute to motor neuron degeneration, with implications for human neurodevelopmental disorders caused by genetic deficiency in Exosome components and genes involved in tRNA maturation.

## RESULTS

### Unspliced pre-tRNAs accumulate in Exosome-deficient cells

To investigate the possible role of the catalytic subunits of mammalian exosome in mature tRNA and/or precursor tRNA (pre-tRNA) surveillance and degradation, we used shRNA to generate stable DIS3 or EXOSC10 knockdown in mouse embryonic stem cells (ESCs) (**Figure S1a**). We performed Northern blots of total RNAs isolated from control and knockdown ESCs using a specific probe against the 5’ exon sequence of tRNA-Tyr-GTA-1 (**Figure 1a, b**). Using Northern probes complementary in sequence to the 5’ exon of intron-containing RNAs allowed us to simultaneously visualize both the unspliced pre-tRNA and spliced form of mature tRNA from each RNA sample. A premature form of tRNA-Tyr-GTA-1 was found to specifically accumulate in the EXOSC10 stable knockdown ESCs, but not in DIS3 stable knockdown cells (**Figure 1b, left panel**). We confirmed this accumulation of unspliced pre-tRNA in EXOSC10-depleted cells using a probe that detects the intron sequence of tRNA-Tyr-GTA-1 (**Figure 1b, right panel**). Similar results were obtained for another intron-containing tRNA, tRNA-Ile-TAT-2 where we observed accumulation of pre-tRNA-Ile-TAT-2 (**Figure S1b, c**). The mature forms of both tRNA-Tyr-GTA-1 and tRNA-Ile-TAT-2 were unchanged in DIS3 or EXOSC10 knockdown cells (**Figures 1b, S1b**). Furthermore, levels of an intronless tRNA, tRNA-Lys-TTT-1, were not affected by EXOSC10 depletion (**Figure 1c**). Taken together, these results support that the Exosome, and particularly EXOSC10, is implicated in pre-tRNA surveillance and decay, whereas the steady state levels of mature tRNAs are largely unaffected by Exosome depletion.

**Figure 1.**
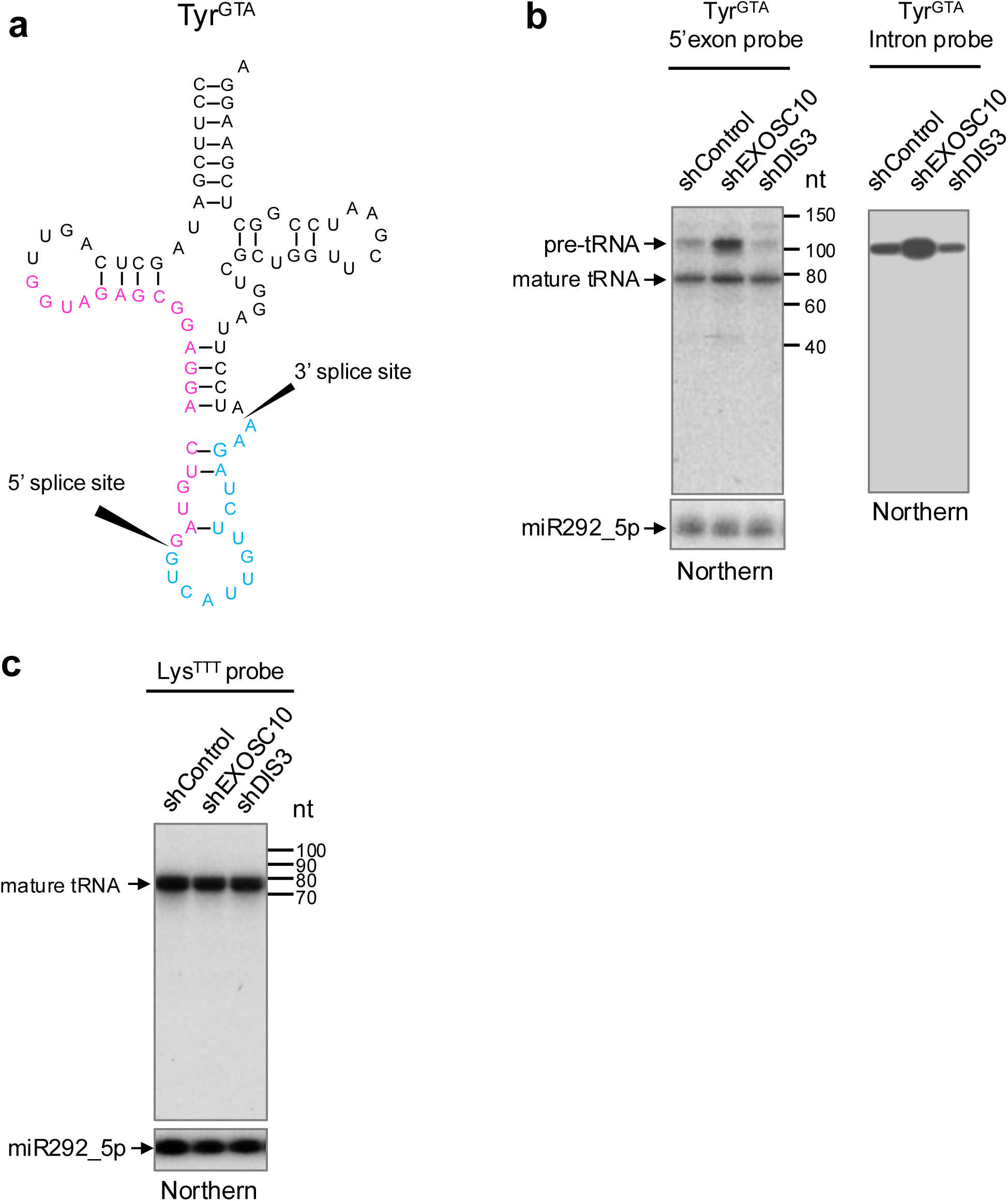
Unspliced precursor tRNA accumulates in EXOSC10-deficient cells. **(a)** Depiction of mouse pre-tRNA-Tyr-GTA-1 sequence and splice sites (solid arrows). The sequence complementary to the 5’exon probe is colored in magenta and the sequence complementary to the intronic probe colored in blue. Probes were used for Northen blot in (b). **(b)** Northern blot analysis of pre-tRNA-Tyr-GTA-1 tRNA performed on RNA samples collected from control, shEXOSC10 and shDIS3 stable knockdown in mouse embryonic stem cells (ESCs). Detection of Pre-tRNA-Tyr-GTA-1 tRNA using the 5’ exon probe (left panel) or reblotted using the intron probe (right panel). The premature form of the intron-including tRNA accumulates in EXOSC10-deficient ESCs. miR292_5p is shown as a loading control. **(c)** tRNA-Lys-TTT-1 detection by northern blotting. miR292_5p is shown as a loading control.

### The Exosome is required for degradation of unspliced pre-tRNAs

To test the direct requirement of EXOSC10 in the degradation of pre-tRNAs, we next used an *in vitro* degradation assay. These experiments were performed using nuclear extracts prepared from control-or EXOSC10 knockdown ESCs. Radiolabeled pre-tRNA substrates were incubated with control or EXOSC10-deficient extracts over a 20-minute time-course. This revealed that both the unspliced yeast pre-tRNA-Phe and the unspliced mouse pre-tRNA-Tyr-GTA-1 were stabilized in the EXOSC10-depleted lysate (**Figure 2a-c**). These results uncover a specific requirement for EXOSC10 in the decay of pre-tRNAs and this likely explains the accumulation of pre-tRNAs observed in cells with compromised EXOSC10 function (**Figure 1**).

**Figure 2.**
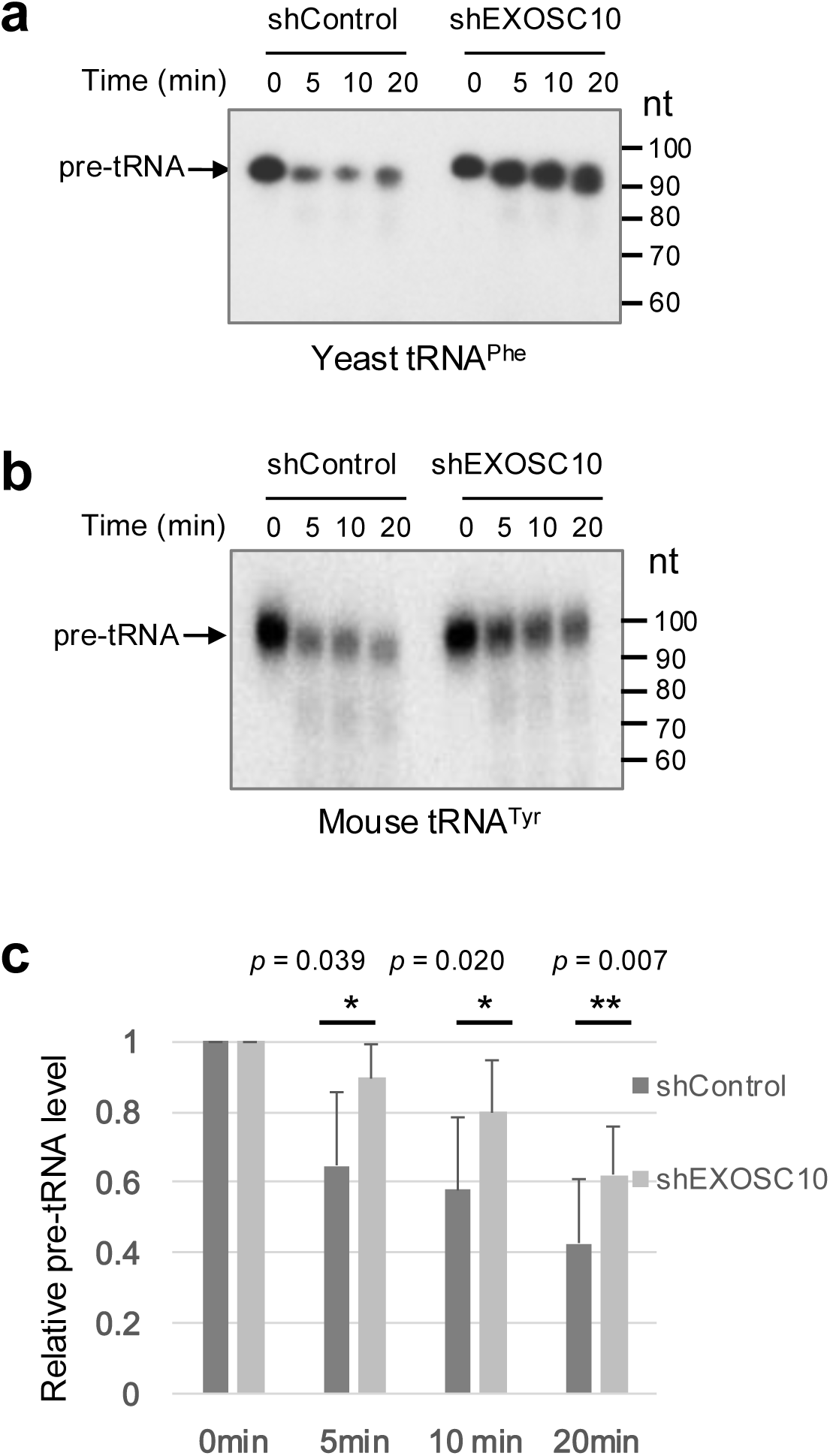
EXOSC10 is required for degradation of unspliced precursor tRNAs. **(a)** *In vitro* pre-tRNA decay assay using radiolabeled yeast pre-tRNA-Phe substrate and nuclear extracts prepared from control and EXOSC10-depleted cells and incubated for the indicated time points **(b)** *In vitro* pre-tRNA decay assay using radiolabeled mouse pre-tRNA-Tyr-GTA substrate and nuclear extracts prepared from control and EXOSC10-depleted cells and incubated for the indicated time points**. (c)** Quantified band intensities from assays with pre-tRNA-Tyr-GTA substrate (as in b). Relative levels of pre-tRNA-Tyr-GTA levels are shown plotted and statistical significance is calculated. Error bars, s.d.; n=3. shControl, control shRNA; shEXOSC10, shRNA to EXOSC10; pre-tRNA, premature tRNA.

### Accumulation of unspliced pre-tRNA impacts embryonic stem cell homeostasis

Next, to explore the possible consequences of pre-tRNA accumulation, we developed a system for overexpression of pre-tRNA-Ile-TAT-2. To accomplish this, we cloned the pre-tRNA sequence into the pLKO.1 lentiviral expression vector. To suppress processing of the pre-tRNA to the spliced form we mutated a single nucleotide within the intron that disrupts a conserved anticodon-intron base-pair that is critical for the splicing step (**Figure S1c**)(Lee and Knapp 1985, Abelson 1992). ESCs were transduced with virus expressing either control vectors or the pre-tRNA expressing vector and stable cell lines were generated. Northern blot of control and pre-tRNA expressing ESCs confirmed a moderate level of pre-tRNA overexpression (**Figure 3a**). To explore the downstream molecular effects of pre-tRNA accumulation, we performed RNA-seq by using poly-A containing mRNA samples from control and pre-tRNA-Ile-TAT-2 expressing ESCs (**Figure 3b**). We identified 1,019 genes with greater than 1.5-fold decreased expression and 157 genes with greater than 1.5-fold increased expression in the pre-tRNA-Ile-TAT overexpressing ESCs compared with controls (**Supplementary Dataset 1)**. Interestingly, gene ontology (GO) analysis of the dysregulated mRNAs revealed that the most significant biological processes associated with the upregulated genes are axonogenesis, neuroprojection, and regulation of cell growth, whereas the most significant biological processes associated with the downregulated genes are chromosome segregation and organelle fission (**Figure 3c**). These results support that a modest level of pre-tRNA overexpression impacts ESC homeostasis, and our unbiased global analysis of the biological processes affected suggests that neuronal pathways might be particularly sensitive to pre-tRNA accumulation.

**Figure 3.**
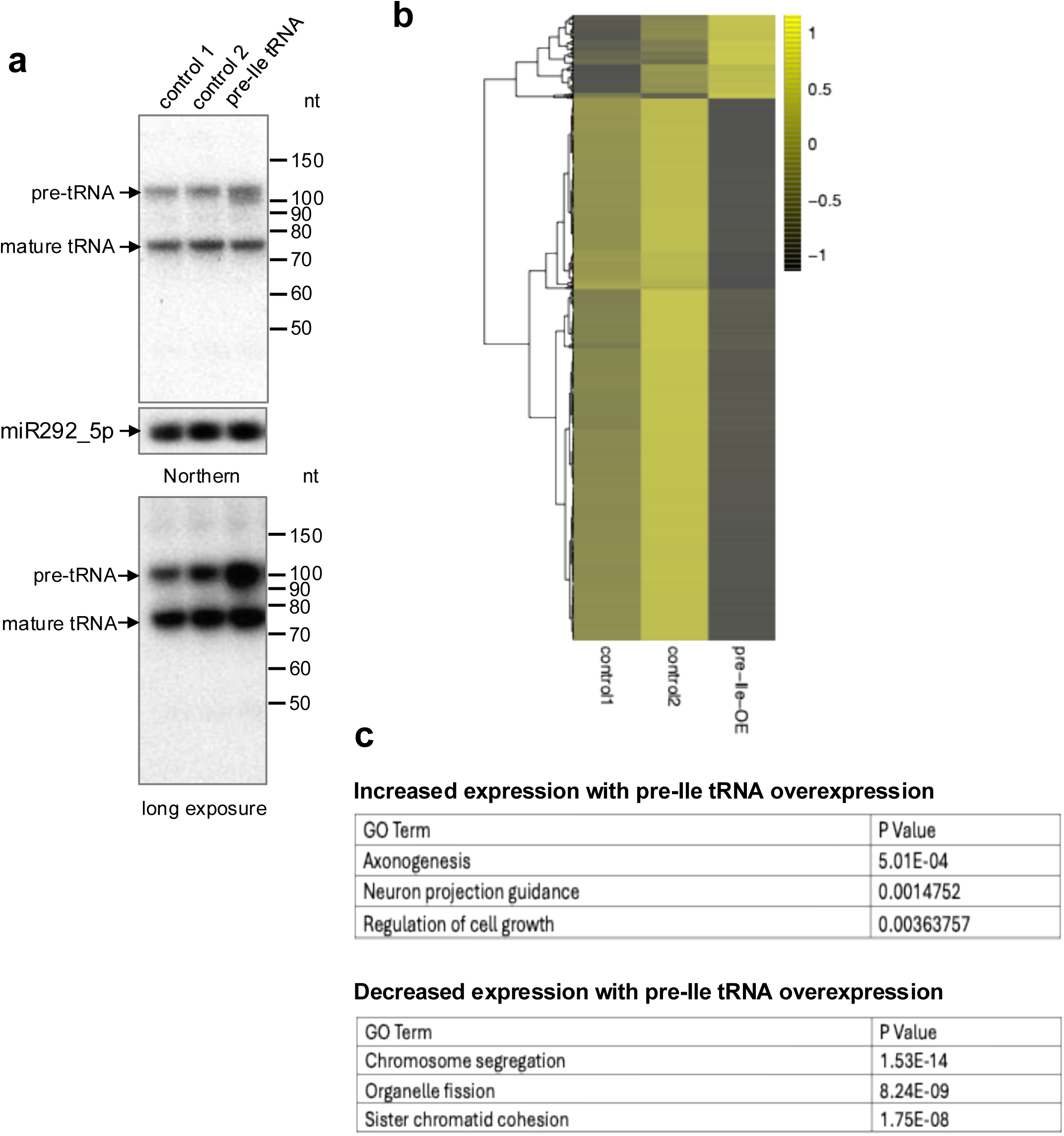
Pathways affected by elevated pre-tRNA expression in ESCs. **(a)** Northern blot of ESCs stably expressing pre-tRNA-Ile-TAT. A longer exposure is shown in the panel below. miR-292_5p is a loading control. Pre-tRNA, premature tRNA; pre-Ile tRNA O/E, premature Ile tRNA overexpression. **(b)** Heatmap showing differentially expressed genes in control 1 ESCs, control 2 ESCs, and ESCs with pre-Ile tRNA O/E. **(C)** Top categories associated with ‘Biological process’ by gene ontology analysis in the up- and downregulated mRNAs caused by pre-Ile tRNA O/E in ESCs. Differentially expressed genes are listed in **Supplementary dataset 1**. pre-Ile tRNA O/E, premature Ile tRNA overexpression; ESC, embryonic stem cell.

### Pre-tRNA inhibits survival of *in vitro* differentiated motor neurons

Considering our RNA-seq results showing the enrichment of neuronal pathways dysregulated in ESCs with pre-tRNA overexpression (**Figure 3**), as well as the established human genetic disorders linking genes involved in tRNA metabolism with neurological diseases, we next sought to test the possible pathological role of pre-tRNAs in neurons. To accomplish this, we used an *in vitro* ESC differentiation assay to examine the effects of pre-tRNA-Ile-TAT overexpression on motor neurons (**Figure S2a**). The high efficiency of motor neuron differentiation was confirmed by immunostaining (**Figure S2b**), and qPCR (**Figure S2c**) analysis of mature motor neuron markers. Apoptotic cells were measured by counting the proportion of cleaved

Caspase3 positive cells in the pan neuronal β tubulin III marker (Tuj1)-positive neurons (**Figure 4a, b**) as well as HB9-positive motor neurons (**Figure 4c, d**). This showed a substantial increase in apoptotic motor neuronal cells caused by pre-tRNA-Ile-TAT overexpression (**Figure 4a-d**). In contrast, pre-tRNA expression did not induce apoptosis in undifferentiated ESCs indicating that the toxic effects might be restricted to certain cell-types including neurons (**Figure S3a-b**).

**Figure 4.**
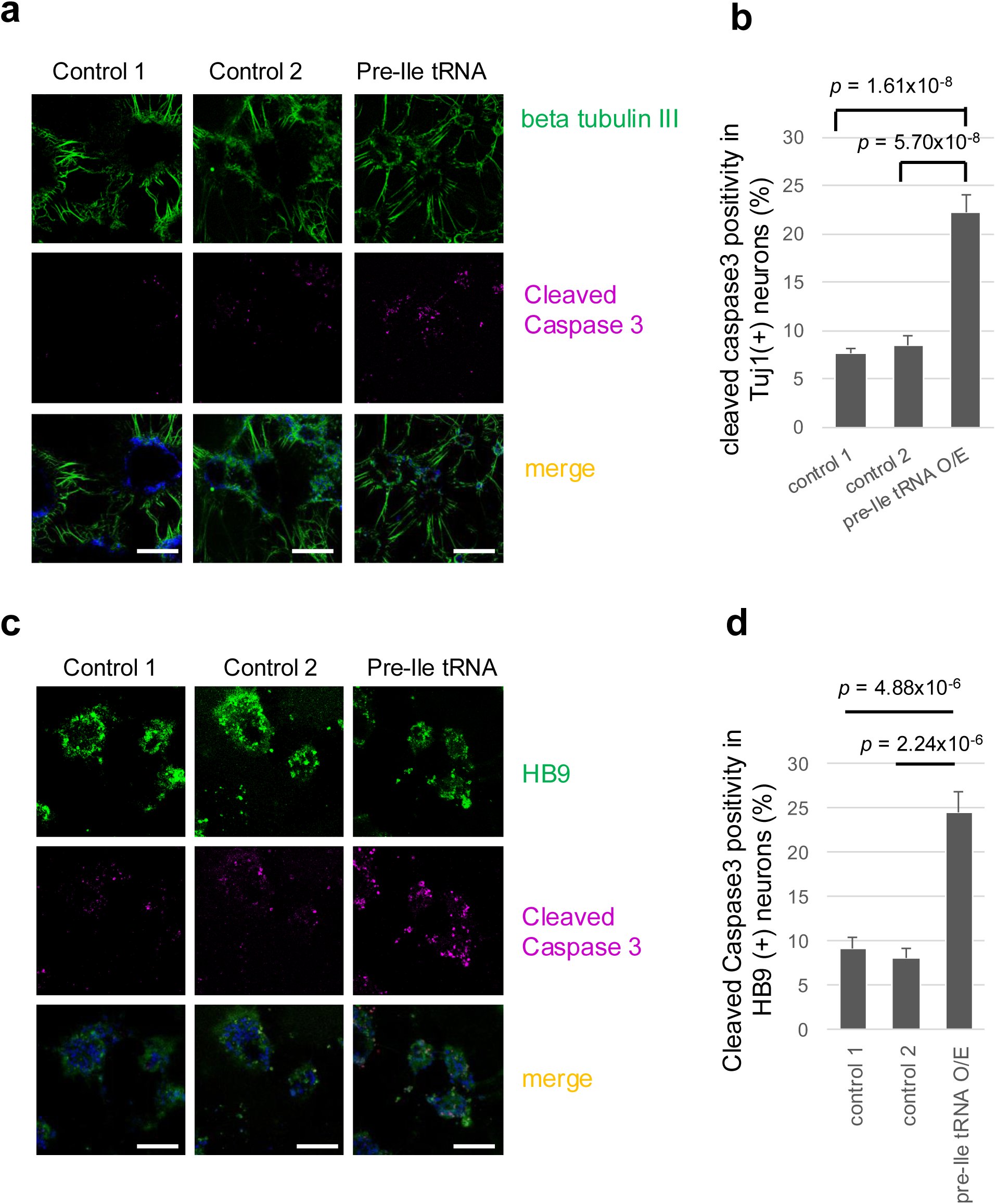
Elevated pre-tRNA expression inhibits neuronal survival. **(a)** Neurons stained with β tubulin III (pan-neuronal marker, green) and cleaved Caspase 3 (cas3, magenta) with DAPI on Day 9 of differentiation. Scale bar, 100 μm. MN, motor neurons. The cleaved Caspase 3 positivity in β tubulin III-positive neurons is quantified and shown in (b). **(b)** Bar graphs showing the proportion of β tubulin III-positive neurons that are cleaved Caspase3 positive in control 1, control 2, and pre-tRNA-Ile tRNA overexpressing cells. Error bars, s.e.m.; 1,157 cells in 11 colonies for the control 1,824 cells in 10 colonies for the control 2, and 866 cells in 22 colonies for pre-tRNA-Ile O/E were counted, respectively. **(c)** Differentiated Motor neurons (MNs) stained with HB9 (green) and cleaved cas3 (magenta) with DAPI on Day 9 of differentiation. Scale bar, 100 μm. The cleaved Caspase3 positivity in HB9-positive mature MNs is shown in (d). **(d)** Bar graphs showing the proportion of HB9-positive neurons that are cleaved Caspase3 positive in control 1, control 2, and pre-tRNA-Ile overexpressing cells. Error bars, s.e.m.; 351 cells in 16 colonies for the control 1,259 cells in 15 colonies for the control 2 and 360 cells in 13 colonies for pre-tRNA-Ile O/E were counted, respectively.

## DISCUSSION

Altogether, our results identify a mammalian pre-tRNA decay pathway involving the EXOSC10 catalytic subunit. Deficiency in this pathway leads to accumulation of unspliced pre-tRNA *in vitro* and in cells, and that elevated pre-tRNA expression induces apoptosis in motor neurons. Our findings provide insight into how Exosome deficiency and pre-tRNA accumulation can contribute to motor neuron degeneration, with implications for human neurodevelopmental disorders caused by genetic deficiency in Exosome components and genes involved in tRNA maturation (Wan, Yourshaw et al. 2012, Boczonadi, Muller et al. 2014, Blaze and Akbarian 2022). Moreover, a recent genetic study in mice demonstrated an essential role of Exosc10 in brain development, with Exosc10 ablation in early cortical progenitors causing massively enhanced apoptosis, reduced neurogenesis, and dysgenesis of cortical layers of the brain (Ulmke, Xie et al. 2021). It will be of interest to examine pre-tRNA levels in this context and explore whether pre-tRNA accumulation might be contributing to these developmental phenotypes in Exosc10-deficient brains.

Importantly, we find that a relatively modest level of pre-tRNA overexpression can specifically induce apoptosis in motor neurons but not in undifferentiated ESCs. However, a previous study found that the 3’ exon fragment (in particular, a 3’ fragment with a 5’-OH group) but not pre-tRNAs (or 5’ exon fragment) could have toxic effects on the viability of CLP1 mutant human fibroblasts and induced neuronal cells (iNeurons)(Schaffer, Eggens et al. 2014). The difference in our observations might be due to cell-type specific effects or different sensitivity of the assays used. These findings provide important new insight into how Exosome deficiency is linked with the pathogenesis of PCH1. An important outstanding question is why premature tRNA is toxic to motor neurons and what are the downstream pathways? Ongoing and future work will help uncover the exact cellular specificity and downstream pathways involved in the toxicity of pre-tRNAs and splicing intermediates. The demonstration that neonatal lethality and motor neuron loss in a mouse model of CLP1 deficiency, could be rescued by P53 loss provides some important insight into this question (Hanada, Weitzer et al. 2013).

## MATERIALS AND METHODS

### Cell culture

Control, shDis3, and shExosc10 stable V6.5 ES cell lines (ESCs), and control, and pre-tRNA-Ile-TAT overexpressing stable V6.5 ESCs were generated and maintained on feeder cells in DMEM supplemented with 15% FBS, 1,000 units/mL mLIF and 2.5 µg/mL puromycin, or on 0.1% gelatin-coated dish in DMEM/F12 and Neurobasal mixture medium supplemented with N-2 Supplement, B27 supplement and 2i (1 µM PD0325901 and 3 µM CHIR99021) with 1,000 units/mL mLIF and 2.5 µg/mL puromycin (Chang, Triboulet et al. 2013). HEK293T cells were maintained in DMEM supplemented with 10% FBS.

### Western blotting

Cells were rapidly frozen in liquid N2 and lysed with lysis buffer (137 mM NaCl, 20 mM Tris-Cl (pH 8.0), 1 mM EDTA, 1% Triton X-100, 10% Glycerol, 1.5 mM MgCl2) including protease inhibitor complete (11697498001, Roche). Proteins were run on 4%–12% Tris-Glycine gels (Invitrogen) and transferred to PVDF membrane (Millipore). Antibodies used were anti-Exosc10 antibody (Abcam, ab50558, 1:1,000), anti-Dis3 antibody (Proteintech, 14689-1-AP, 1:1,000) and anti-β actin antibody (Abcam, ab75186, 1:2,000).

### RNA extraction and Northern blot

Total RNA was extracted using Trizol (Invitrogen) according to the manufacturer’s instruction. Northern blot was performed as previously described using 5 or 10 µg of RNA. Specific probes for the Tyr 5’ exon or the intron of *M. musculus* tRNA-Tyr-GTA-1 (mouse July 2007 (mm9) genome assembly), and for the Ile 5’ exon against M. musculus tRNA-Ile-TAT-2, and for the Lys tRNA against *M. musculus* tRNA-Lys-TTT-1 were purchased from IDT. The probe sequences are listed in **Supplementary Table 1**.

### Stable knockdown and pre-tRNA overexpression in ESCs

Stable V6.5 ESCs were generated expressing MISSION short hairpin RNA (shRNA) including control shRNA, Exosc10 shRNA, and Dis3 shRNA. To construct the stable pre-tRNA-Ile-TAT overexpressing cell lines, pLKO.1-puro vector (Addgene) was digested with AgeI and EcoRI and tRNA-annealed oligos (sense: 5’-CCGGGCTCCAGTGGCGCAATCGGTTAGCGCGCGGTACTTATACAGCAGTACATGCAGA<u>C</u>CAATGCCG AGGTTGTGAGTTCGAGCCTCACCTGGAGCATTTTTG-3’, antisense: 5’-AATTCAAAAATGCTCCAGGTGAGGCTCGAACTCACAACCTCGGCATTG<u>G</u>TCTGCATGTACTGCTGTATA AGTACCGCGCGCTAACCGATTGCGCCACTGGAGC-3’), were inserted. Sequence-verified plasmids were transfected together with pLP1, pLP2, and VSVG vectors, into HEK293T cells for lentivirus production. Note that an underlined base of the conserved anticodon-intron base pair in the pre-Ile-tRNA sequence was mutated to suppress splicing (**Figure S1c)**. The original template pLKO.1-TRC vector including multiple cloning sites was used as control 1 vector. Oligos (sense: 5’-CCGGTCCGCAGGTATGCACGCGTGTTTTTG-3’, antisense: 5’-AATTCAAAAACACGCGTGCATACCTGCGGA-3’) were used for the construction of control 2 vector, in which the multiple cloning sites were removed and ‘TTTTT’ stop signal of transcription was added. After identifying the optimal titers, the virus particles were used to infect V6.5 ESCs. Each stable cell line was established by puromycin (2.5 µg/ml) selection.

### *In vitro* pre-tRNA degradation assay

Nuclear extracts were prepared as described in previous articles and finally dialyzed against a buffer including 30mM Tris-HCl [pH7.4], 100mM KCl, 5mM MgCl_2_, 10% v/v glycerol, 1mM DTT and 0.1mM AEBSF serine protease inhibitor (Sigma A8456) by using Slide-A-Lyzer MINI Dialysis Devices, 3.5K MWCO (Thermo #69552) for 2h at 4° C. The premature yeast Phe^GAA^ tRNA substrate was synthesized by PCR using *S. cerevisiae* genomic DNA, 5’ primer (5’-AATTTAATACGACTCACTATAGGGGATTTAGCTCAGTTGGG-3’) and 3’ primer (5’-<u>TGGTGG</u>GAATTCTGTGGATCGAAC-3’). The RNA template was prepared using the T7 MEGAshortscript kit (Ambion) with 450Ci/mmol UTP, [α-32P] (3,000Ci/mmol EasyTide, Perkin Elmer, BLU507H). The premature tRNA was dissolved at 1 µM in the buffer consisting of 30mM HEPES-KOH [pH 7.3], 2mM MgCl_2_ and 100mM KCl, followed by incubation with the indicated nuclear extracts then resolved using denaturing TBE urea PAGE and radiolabeled RNA visualized by autoradiography.

### RNA-sequencing and data analysis

For RNA sequencing analysis of ESCs, 2 µg of RNA was purified with fragmentation of poly-A containing mRNA, followed by the TruSeq Stranded mRNA Sample Preparation Low Sample protocol (Illumina). RNA-seq was carried out by using the NextSeq 500 (Illumina). Bowtie software (http://bowtie-bio.sourceforge.net/bowtie2/index.shtml) was used for alignment of sequencing reads to mRNA transcriptome (mm9), and a self-developed PERL program was used to calculate the expression level of individual genes. We excluded low abundance mRNAs with an average read number of less than 5 in control 1, control 2 and pre-Ile overexpressing) (O/E) cells. RNAs were selected based on more than ≧1.5-fold increase and decrease in pre-Ile-tRNA O/E ESCs compared with both control ESC samples. Cluster 3.0 and Java Treeview software (http://bonsai.hgc.jp/~mdehoon/software/cluster/) was used for the heatmap analysis. The gene ontology analysis was carried out by using ToppGene software (https://toppgene.cchmc.org).

### Motor neuron differentiation

Mouse ESCs were differentiated into motor neurons by adding differentiating factors shown in **Figure S2a** (Wichterle, Lieberam et al. 2002, Wichterle and Peljto 2008, Maeda, Harris et al. 2014). Briefly, mouse ESCs were dissociated and plated on the Ultra-low attachment culture dish (Sigma-Aldrich), cultured in ADFNK medium consisting of Advanced DMEM/F12 (Gibco): Neurobasal medium (Gibco) 1:1, PC/SM, L-Glutamine, 0.1 mM β-mercaptoethanol, 10% Knockout serum replacement (Gibco). On Day 2, 1 µM of RA (Sigma) was supplemented to the medium. On Day 3, 200 ng/mL of Sonic hedgehog (Shh, R&D system) and on Day 5, 10 ng/mL of GDNF (R&D system). On Day 6, differentiated cells in the form of embryonic bodies (EB) were collected in Trizol, used for RNA collection. For the immunostaining, the EB-formed spheres were embedded in Tissue-Tek OCT compound after fixation using 4% paraformaldehyde (PFA) and equilibration with 30% sucrose solution. 30 µm thick sections were used for immunocytochemistry. For the single cell staining, EB-formed differentiated cells were dissociated using Accumax (Millipore) and 3×10^5^ cells /cm^2^ were plated on 300 µg/mL of Matrigel Matrix Basement Membrane (BD Bioscience) on glass bottom dish (Mat Tek) coated with 100 µg/mL poly-DL-ornithine and 2 µg/mL of laminin (Millipore) on Day 6. The cells were cultured in the ADFNK medium including RA, Shh, GDNF, 10 ng/mL BDNF (R&D system), 10 ng/mL CNTF (R&D system) and 10 ng/mL NT-3 (R&D system) Day 6 through Day 9 (Maeda, Harris et al. 2014). On Day 9, cells were fixed by using 4% PFA and 2% glutaraldehyde, followed by the immunocytochemistry.

### Reverse transcription and quantitative PCR

Total RNA extracted from V6.5 ESC and at Day 6 of *in vitro* differentiation was reverse transcribed using random hexamers and SuperScript III according to the manufacturer’s instruction (Invitrogen). Each mRNA level was analyzed by qPCR using Fast SYBR Green Master Mix and StepOnePlus Real-Time PCR System (Applied Biosystems). The sequence of specific primers used for qPCR are listed in **Supplementary Table 1**.

### Cytospin funneling

ESCs were suspended in PBS after trypsinization, followed by the spin coating with 1,500 rpm, 3 min onto the glass slides by using Shandon EZ Single Cytofunnel. After fixation with 4% PFA, immunocytochemistry was performed.

### Immunocytochemistry

Fixed samples were permeabilized with PBS including 0.1% Triton X at RT for 20 min and blocked with PBS including 10%FBS and 0.1% Triton X at 4C, overnight. Cells were incubated at RT for 1 hour with primary antibodies β III tubulin (rabbit polyclonal, Abcam, 1:100; mouse monoclonal, Thermo Scientific MA1-118, 1:100), HB9 (mouse monoclonal, DSHB 81.5C10, 1:10), Islet-1 (mouse monoclonal, Islet-1/2 homeobox, DSHB 39.4D5, 1:10), Olig2 (mouse monoclonal, Millipore MABN50, 1:100), PAX6 (mouse monoclonal, DSHB, 1:10) and cleaved Caspase 3 (rabbit polyclonal, Cell Signaling, 1:100). The cells were then washed with PBS and incubated with the secondary antibodies Goat Anti-Mouse IgG H&L (Alexa Fluor 488, 1:100), Goat Anti-Rabbit IgG H&L (Alexa Fluor 488, 1:100), Goat Anti-Mouse IgG H&L (Alexa Fluor 594, 1:100) and Goat Anti-Rabbit IgG H&L (Alexa Fluor 594, 1:100) for 1 hour at room temperature. The coverslips were mounted with Vectashield with DAPI (Vector Laboratories, Burlingame, CA) and observed with Leica SP2 scanning confocal microscope (Leica, Wetzlar, Germany).

### Statistical Analysis

For all quantified data, mean ± s.d. or mean ± s.e.m. are presented. Statistical comparisons were made using the Student’s t test. The p values less than 0.05 were considered significant in this study.

## ACKNOWLEDGMENTS

We are grateful to Drs. Akiko and Tadanori Mammoto in the Vascular Biology Program at Boston Children’s Hospital for their help with microscopy. The monoclonal antibodies, anti-HB9 and anti-Islet-1 developed by HHMI/Columbia University and anti-PAX6 developed by Tokyo Institute of Technology, were obtained from the Developmental Studies Hybridoma Bank (DSHB), created by the of the National Institute of Child Health and Human Development (NICHD) of the NIH and maintained at The University of Iowa. Y.N. was supported by the Japanese Society for The Promotion of Science (JSPS)(a postdoctoral fellowship for research abroad and 21K07281), the Uehara Memorial Foundation, the Waksman Foundation of Japan Inc., and the KANAE foundation for the promotion of medical science.

## AUTHOR CONTRIBUTIONS

Y.N. performed all experiments. Y.N. and R.I.G. designed the study, interpreted the results, and wrote the manuscript. X.Y. performed bioinformatics analysis. All authors declare no competing financial interests. Correspondence and requests for materials should be

## FIGURE LEGENDS

**Figure S1.**
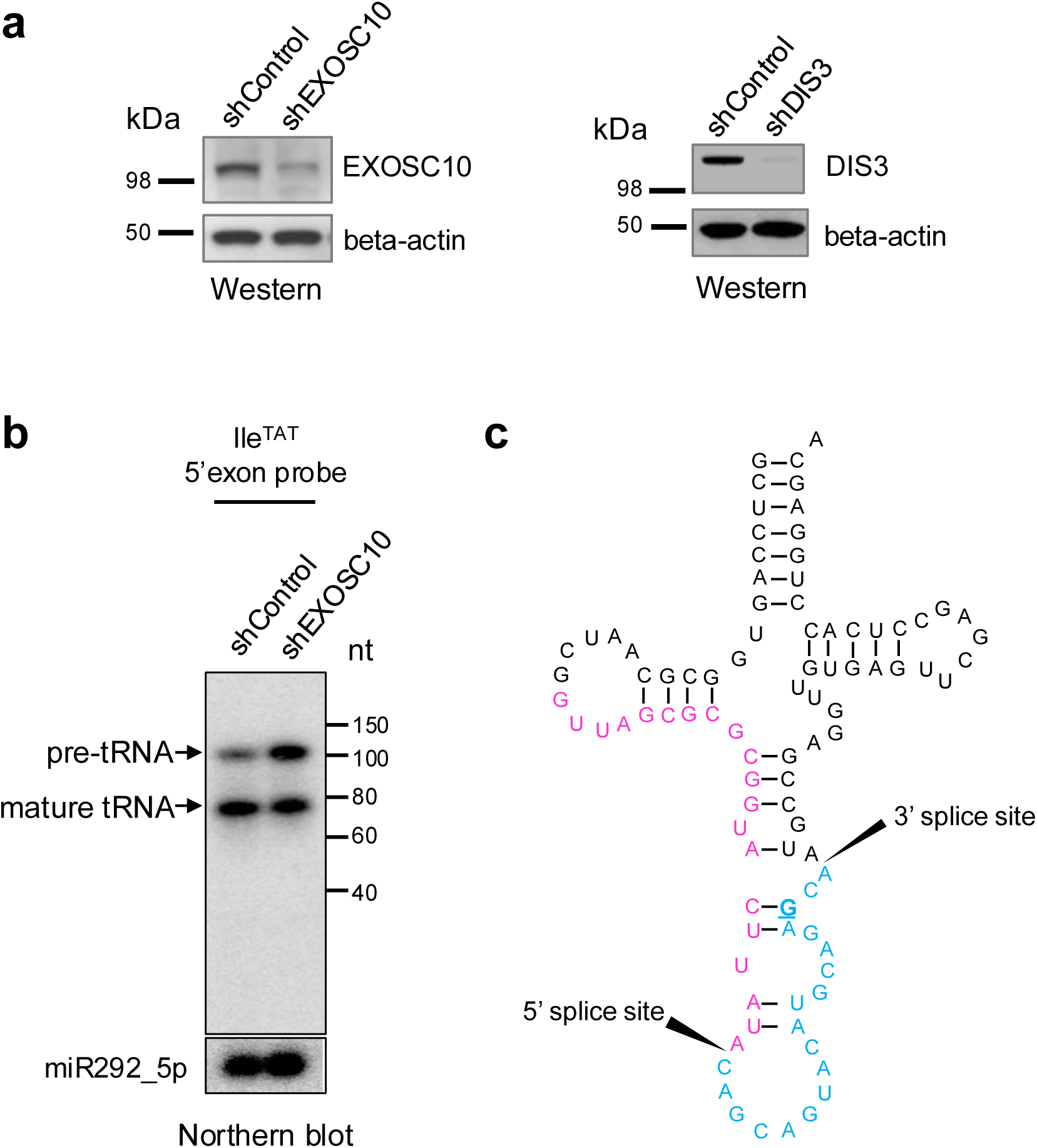
Exosome knockdown and pre-tRNA accumulation. **(a)** Western blot showing stable EXOSC10-and DIS3 knockdown in mouse embryonic stem cells (ESCs). **(b)** Detection of Pre-tRNA-Ile-TAT-2 by Northern blotting using the 5’exon probe in samples from control or EXOSC10 stable knockdown ESCs. Pre-tRNA-Ile-TAT-2 accumulates in EXOSC10-deficient ESCs. miR292_5p Northen blot is shown as a loading control. **(c)** Depiction Pre-tRNA-Ile-TAT-2 sequence. The sequence complementary to the 5’exon probe is colored in magenta and the intronic sequence colored in blue. Splice sites are indicated by solid arrows. Underlined and bold ‘G’ is a critical base for tRNA splicing and is mutated to ‘C’ for pre-tRNA-Ile overexpression experiments shown in **Figures 3 and 4**.

**Figure S2.**
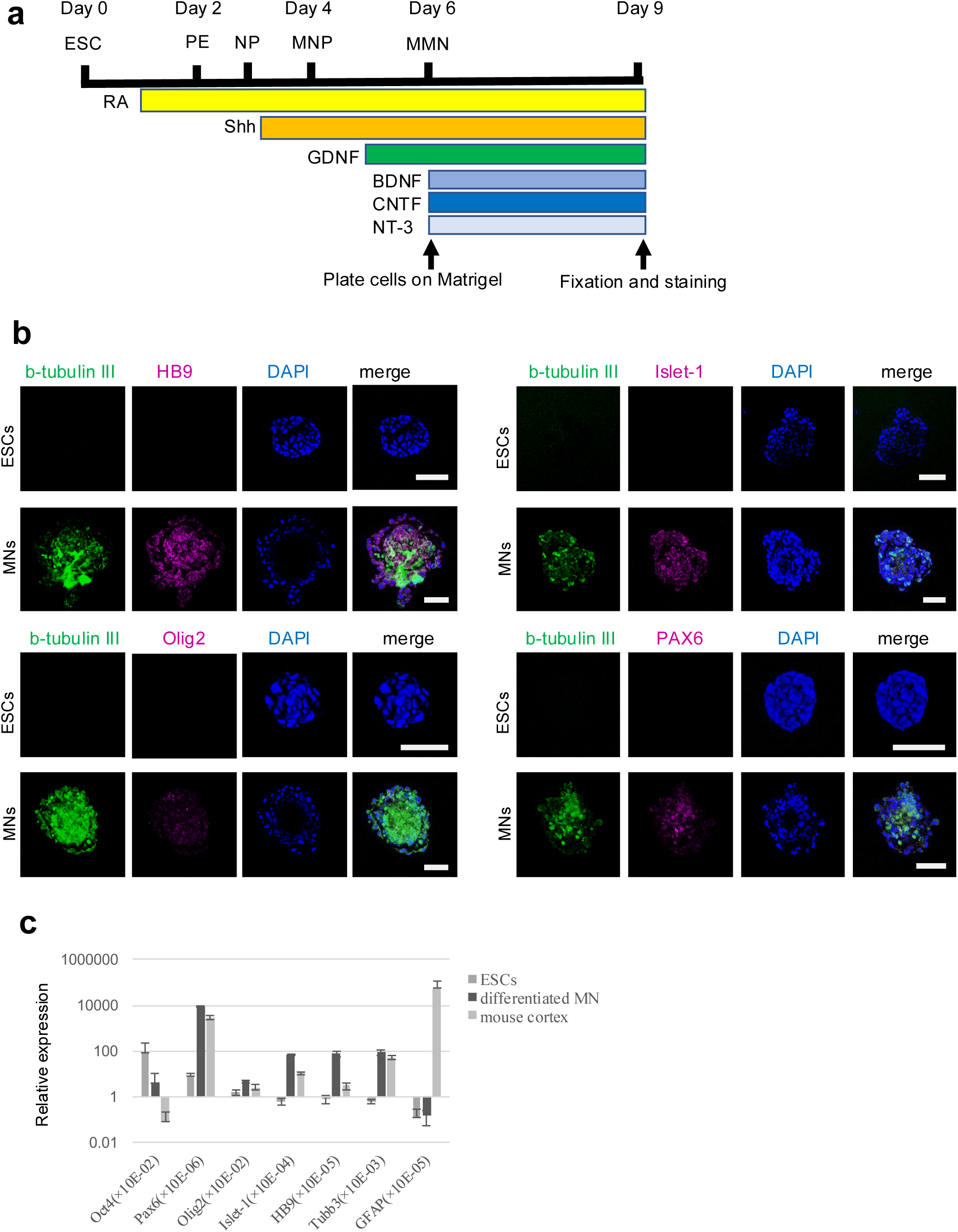
Differentiation of mouse ESCs into motor neurons. **(a)** Scheme of the *in vitro* differentiation strategy. Chemicals and neurotrophic factors added on each day are described below the time course. ESC, embryonic stem cells; PE, primitive ectoderm; NP, neural plate; MNP, motor neuron progenitor; MMN, mature motor neuron. **(b)** Immunocytochemistry of the spheres of mature motor neurons at Day 6. Note that mature motor neuron-specific markers, HB9 and Islet-1, were expressed at Day 6. Scale bars, 50 μm. **(c)** q.RT-PCR analysis of RNA collected from ESCs, differentiated motor neurons (MN) harvested at Day 6, and adult mouse cortex (positive control). The ratio of each of the indicated differentiation marker genes relative to the level of GAPDH is presented. The expression levels relative to GAPDH were 85.8, 4.0 and 0.077 ×10^-2^ for Oct4, 9.1, 8675.4 and 2812.6 ×10^-6^ for Pax6, 1.6, 4.9 and 2.6 ×10^-2^ for Olig2, 0.6, 71.9 and 11.0 ×10^-4^ for Islet-1, 0.8, 77.9 and 3.0 ×10^-5^ for HB9, 0.6, 85.4 and 49.9 ×10^-3^ for tubulin-III (Tubb3) and 0.1, 0.1 and 56792.6 ×10^-5^ for GFAP, respectively. Error bars are +/-s.d., n=3.

**Figure S3.**
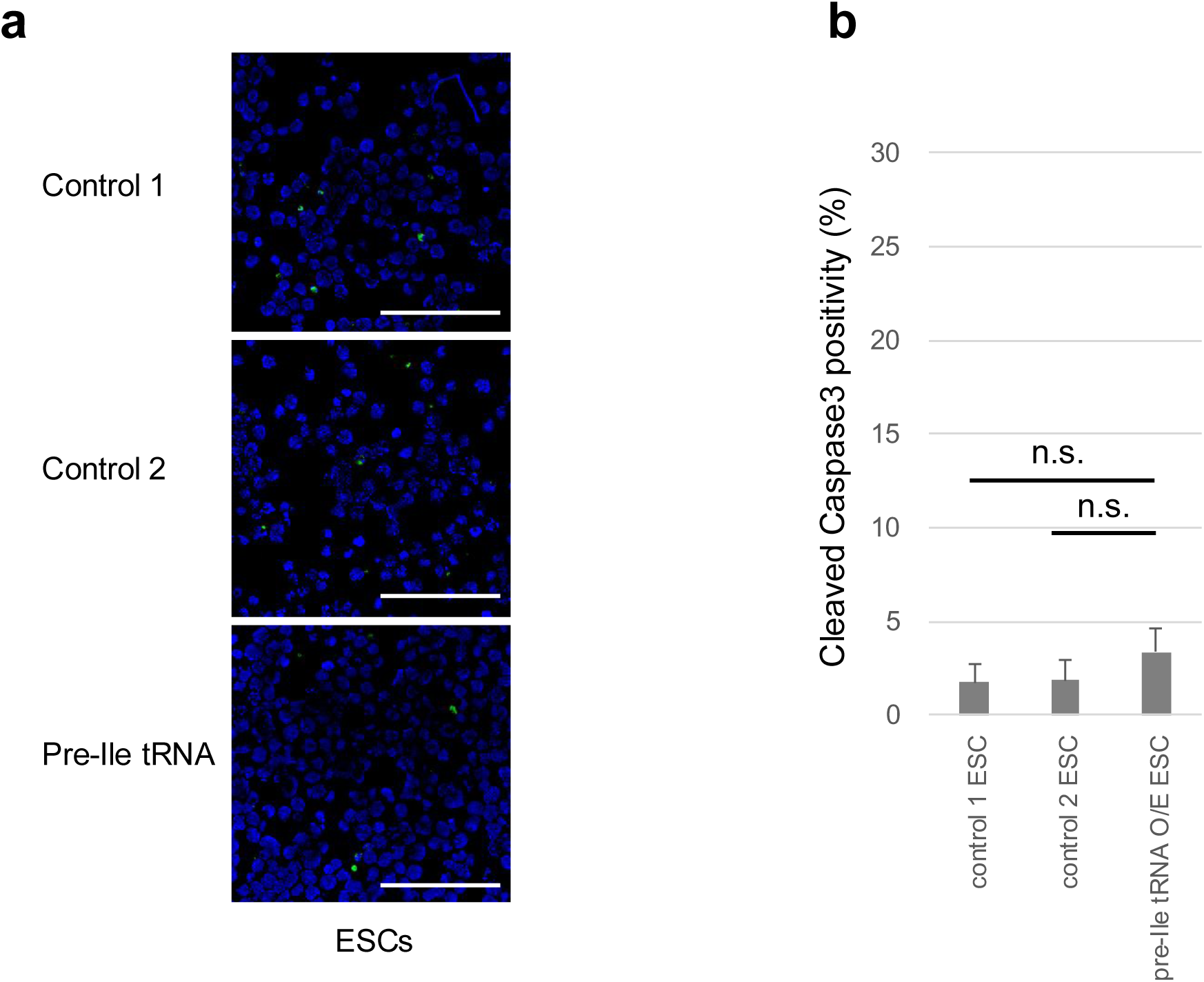
Effect of pre-tRNA-Ile overexpression in ESCs. **(a)** ESCs stained with α-cleaved Caspase-3 (green) and with DAPI by the cytospin method. Scale bar, 100 μm. **(b)** The cleaved Caspase3 positivity in DAPI-positive ESC is shown in bar graphs on the right. The percentages of α-cleaved Caspase-3 cells in DAPI-positive ESCs are 1.7±1.0% in control 1; 1.9±1.0% in control 2; and 3.3±1.3% in pre-tRNA-Ile overexpressing cells, respectively. Error bars, s.e.m. n=4. A total of 424 cells for the control 1, 608 cells for the control 2, and 684 cells for pre-tRNA-Ile O/E were counted, respectively. n.s.= not significant.

**Supplementary Table 1.**
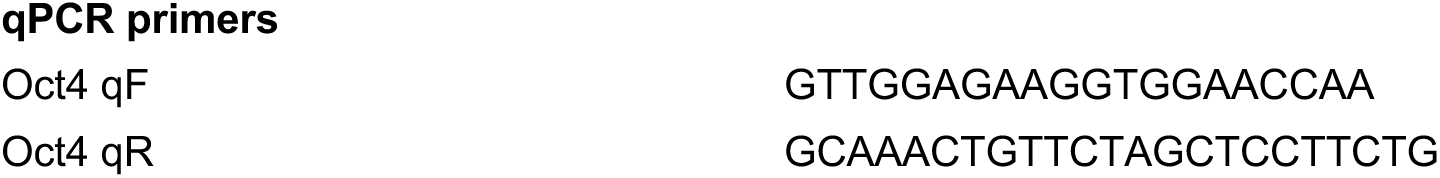
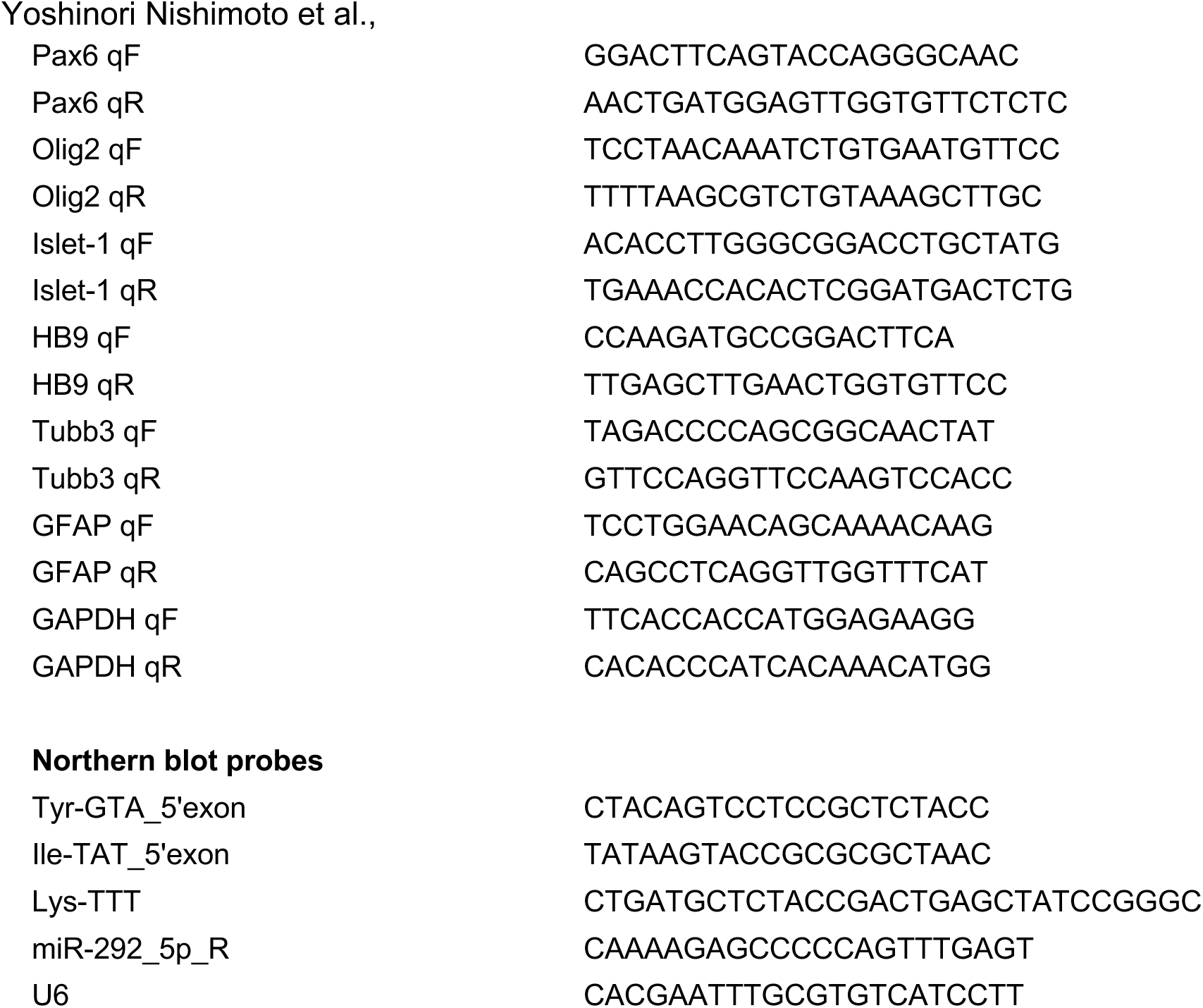
DNA sequence of PCR primers and northern blot probes used in this study

**Supplementary Dataset 1. List of differentially expressed mRNAs in ESCs with elevated pre-tRNA-Ile expression.** The genes are excluded if average read numbers in control 1, control 2, and pre-tRNA-Ile overexpression (OE) are less than 5. Genes with 1.5-fold increased or decreased expression compared to both control 1 and 2 are listed. ESC, embryonic stem cell; pre-tRNA-Ile O/E, precursor tRNA Ile overexpression.

